## Supplementary_Figures for "High frequency body site translocation of nosocomial *Pseudomonas aeruginosa*"

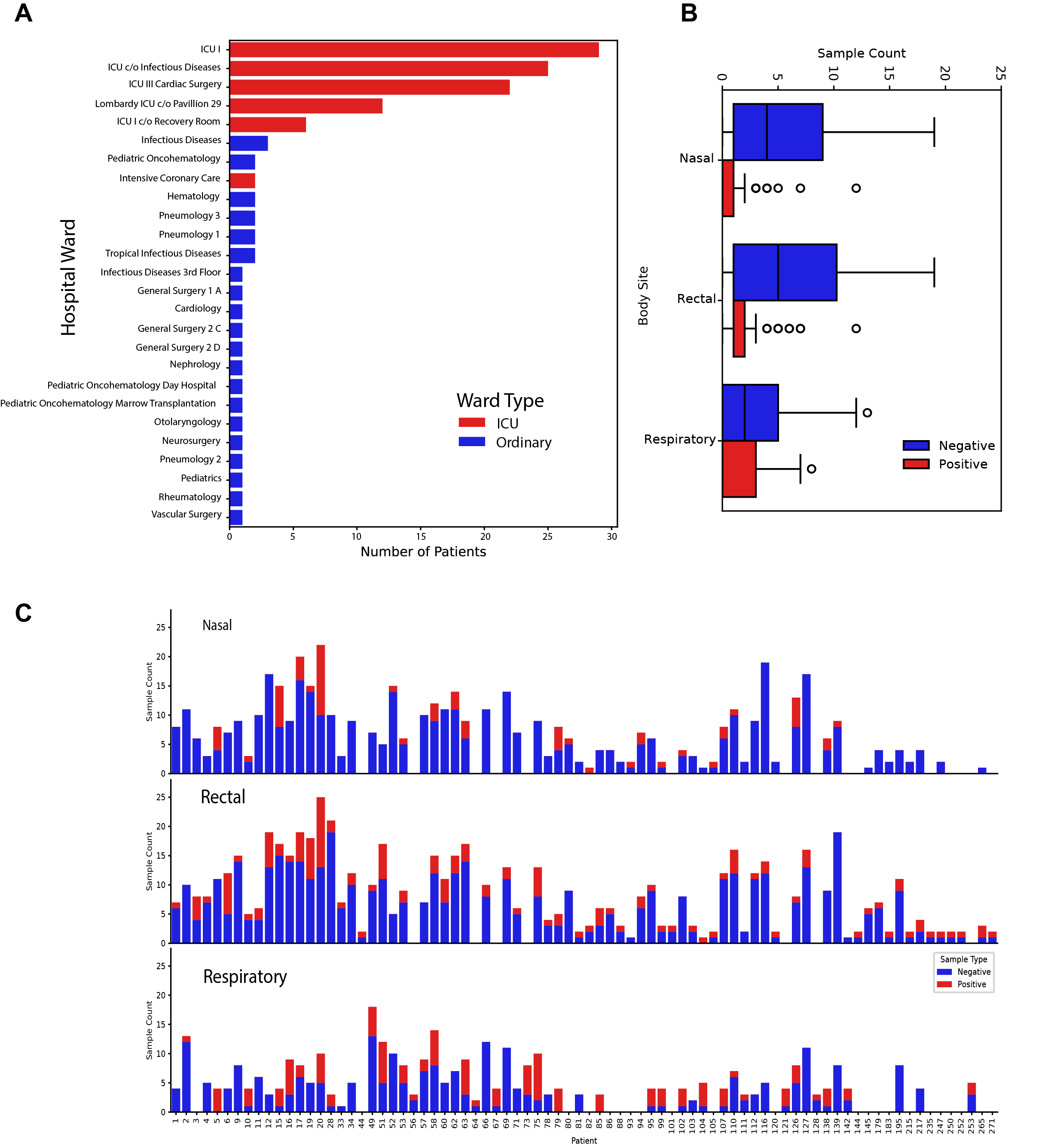

Extended Data Fig.1: Within hospital *P. aeruginosa* distribution across wards and pathogen presence by sample type

A, The number of patients residing in each ward coloured by ward type. Intensive care units (ICU) in red, Ordinary wards in blue. B and C, The number of positive (red) and negative (blue) samples per patient within each body site.

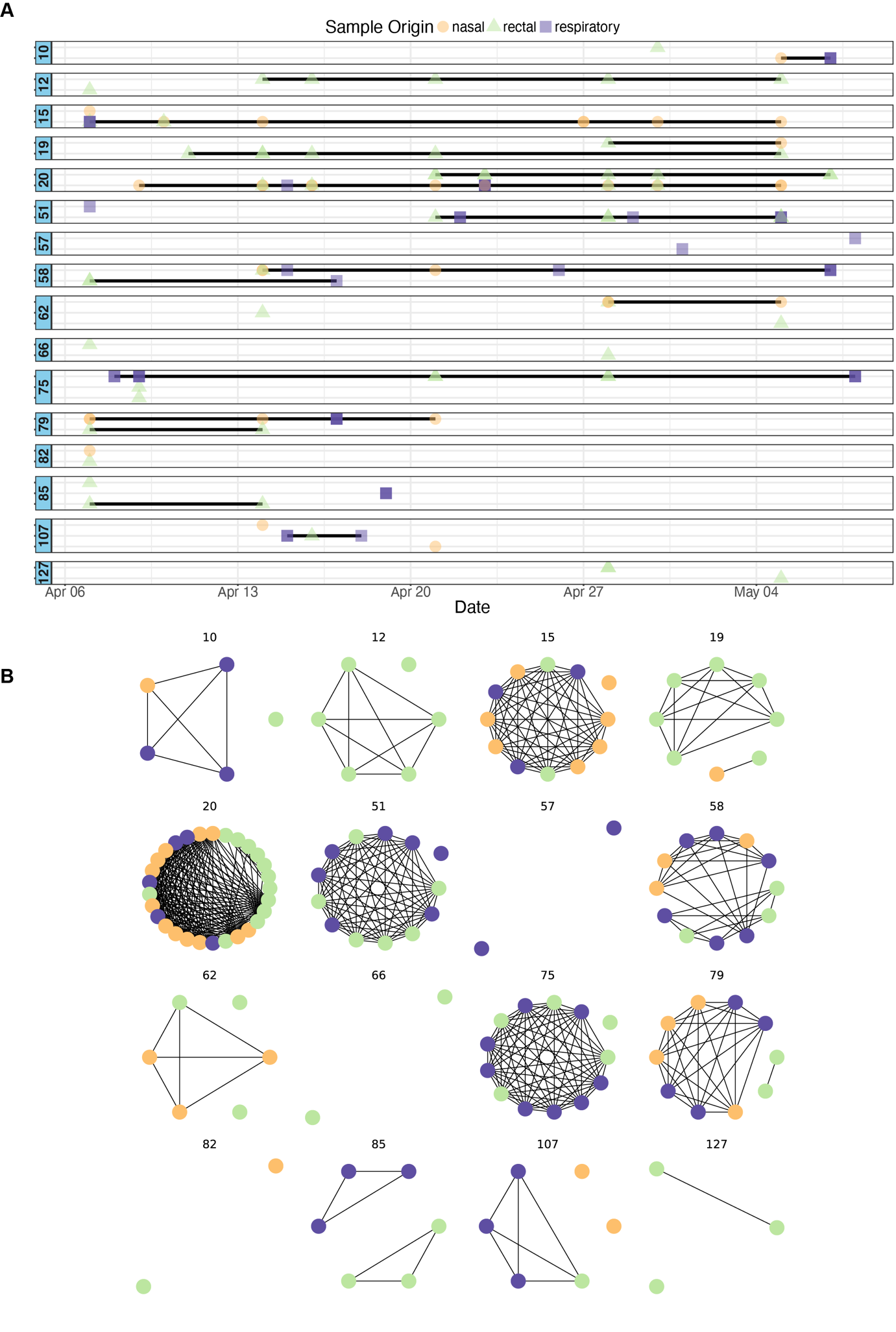

Extended Data Fig.2: Sixteen patients with multiple clones. A, Each point indicates a sample positive for *P. aeruginosa* coloured by sample type. Respiratory samples were coloured purple, rectal samples were coloured green, and nasal samples were coloured yellow. Where clones persisted in separate time point, a black line is used to join the points. B, Network graphs are labelled with patient number. Each node represents a sample (green = rectal, purple = respiratory, yellow = nasal) and each edge shows a difference of less than 100 SNP differences between samples.

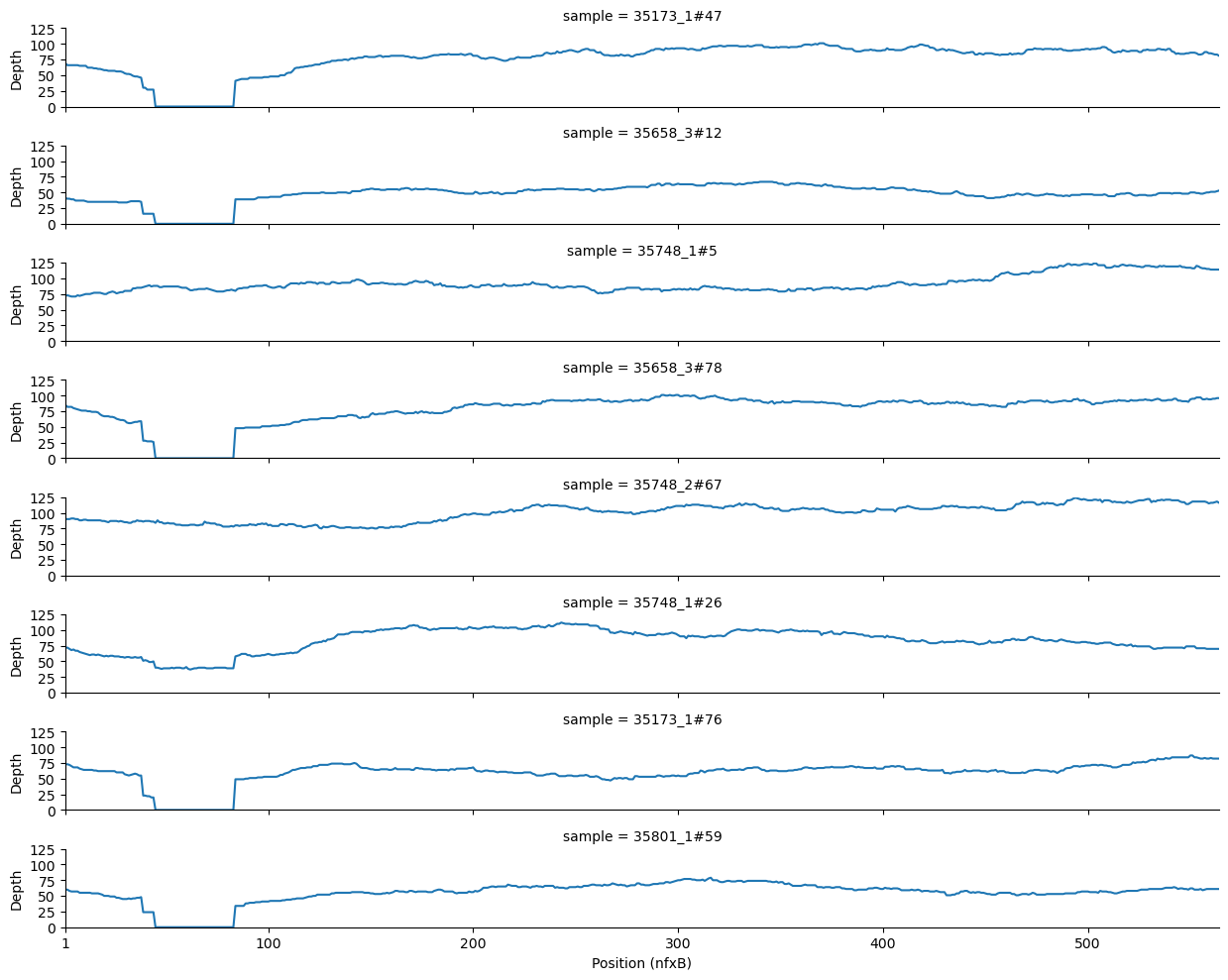

Extended Data Fig.3: Missing coverage in *nfxB* genes within patient 20 clone B.

Extended Data Table.1: *P. aeruginosa* sample metadata

Included within the table are ENA accession numbers, sample types, sequence type, and fastbaps clusters.
